# PIM Kinases Alter Mitochondrial Dynamics and Chemosensitivity in Lung Cancer

**DOI:** 10.1101/863811

**Authors:** Shailender S. Chauhan, Rachel K. Toth, Corbin C. Jensen, Andrea L. Casillas, David F. Kashatus, Noel A. Warfel

## Abstract

Resistance to chemotherapy represents a major obstacle to the successful treatment of non-small cell lung cancer (NSCLC). The goal of this study was to determine how PIM kinases impact mitochondrial dynamics, ROS production, and response to chemotherapy in lung cancer. Live cell imaging and microscopy were used to determine the effect of PIM loss or inhibition on mitochondrial phenotype and ROS. Inhibition of PIM kinases caused excessive mitochondrial fission and significant upregulation of mitochondrial superoxide, increasing intercellular ROS. Mechanistically, we define a signaling axis linking PIM1 to Drp1 and mitochondrial fission in lung cancer. PIM inhibition significantly increased the protein levels and mitochondrial localization of Drp1, causing marked fragmentation of mitochondria. An inverse correlation between PIM1 and Drp1 was confirmed in NSCLC patient samples. Inhibition of PIM sensitized NSCLC to chemotherapy and produced a synergistic anti-tumor response *in vitro* and *in vivo*. Immunohistochemistry and transmission electron microscopy verified that PIM inhibitors promote mitochondrial fission and apoptosis *in vivo*. These data improve our knowledge about how PIM1 regulates mitochondria and provide justification for combining PIM inhibition with chemotherapy in NSCLC.

## Introduction

Lung cancer is the second most commonly diagnosed type of cancer and the leading cause of cancer-related mortality worldwide. More than two-thirds of lung cancer patients are diagnosed at an advance stage (III–IV), and intrinsic and/or acquired resistance to treatment represent major obstacles to the successful treatment of patients with advanced disease (1,2). As compared to other types of lung cancer, non-small cell lung carcinomas (NSCLC) is less prone to undergo spontaneous and treatment-induced apoptosis (3), suggesting that deficiencies in the apoptotic process may be responsible for their and/or acquired resistance to chemotherapy (4).

Cumulative evidence has demonstrated that an imbalance of mitochondrial fission and fusion is common in cancer (5). Mitochondria exist as a dynamic network that is constantly undergoing fusion (elongation) and fission (fragmentation). Mitochondrial fusion results in a tubular mitochondrial network that serves to counteract metabolic insults, maintain cellular integrity, and provide protection against cell death (6,7). In contrast, mitochondrial fission creates small and fragmented mitochondria, which can have both pro- and anti-tumor effects depending on the cellular context (8). Mitochondrial fission is required for cell division and has been shown to positively regulate cell proliferation in cancer cells (9). However, in response to apoptotic stimuli and cellular stress, too much fission generates excessive reactive oxygen species (ROS) and is a necessary event for the initiation of apoptosis (10). Mitochondrial fusion is associated with chemoresistance in several cancer types, including lung cancer (7,11). Moreover, excessive mitochondrial fragmentation is associated with chemosensitive cancer cells in response to platinum-based therapy, whereas chemoresistant cancer cells retain a more fused mitochondrial network (12). These data demonstrate that mitochondrial fusion is an important event in the acquisition of chemoresistance.

An essential step for mitochondrial membrane fission is the recruitment of dynamin-related protein 1 (Drp1) to mitochondria. Drp1 is recruited from the cytoplasm to the mitochondrial outer membrane (13,14), where it colocalizes with receptors (15). Drp1 protein oligomerize to form a ring around the mitochondria that constricts and separates the mitochondrial membrane. Previous reports indicate that downregulation of Drp1 limits mitochondrial fragmentation, cytochrome c release, caspase activation and apoptosis (16), and reduced levels of Drp1 have been reported in many cancers, including NSCLC. Adenocarcinomic alveolar epithelial cells, a model of NSCLC, display decreased Drp1 protein expression, which promotes elongation of mitochondrial phenotypes and limits fission while inhibiting the downstream processes of apoptotic activation (17). These observations suggest that enhancing mitochondrial fission is a promising approach to inducing apoptosis in lung cancer (18).

The *<u>P</u>roviral Integration site for Moloney murine leukemia virus* (PIM) kinases are crucial regulators of cell survival and proliferation, and their expression is associated with poor prognosis in several types of cancer (19). A majority of the research on PIM1 has focused on cancers of hematopoietic, prostate or breast origin, and the mechanisms by which PIM1 promotes therapeutic resistance in lung cancer are poorly understood. Our recent work demonstrated that PIM inhibitors cause a marked increase in reactive oxygen species (ROS), which is crucial for their cytotoxic effects toward cancer cells (20). However, the mechanism by which PIM inhibition produces excessive ROS is not well understood. Here, we investigate the consequence of altered PIM kinase expression or activity on mitochondrial dynamics, ROS and therapeutic resistance in lung cancer.

## Results

### High PIM1 is predictive of poor clinical outcome in NSCLC

To study the clinical significance of PIM1 expression in lung cancer, patient samples and publicly available TCGA datasets of human lung cancer cases were analyzed. Immunohistochemical staining of an NSCLC tissue microarray (TMA) comprising normal lung tissue and patient tumors ranging from stage I – III (n = 100 cores) revealed that PIM1 levels were significantly higher levels in all lung cancer cases relative to normal tissue. Patients with stage II and III had significantly higher PIM1 expression than those with stage I (Fig 1A), and the average PIM1 expression in stage III was two-fold higher than that in stage II (Fig 1B). Moreover, lung cancer patients with high PIM1 expression had significantly worse survival than those with low PIM1 expression. The median survival time of lung adenocarcinoma patients with high PIM1 expression was significantly shorter than that of patients with low expression at each stage [stage I: 111 vs. 68 mo, stage II: 66 vs. 21 mo, and stage III: 34 vs. 23 mo] (Fig 1C). Notably, patients with high PIM1 displayed significantly worse response to chemotherapy than patients with low PIM1 [stage I: 40 vs. 5 mo, and stage II: 16 vs. 5 mo] (Fig 1D). These, findings suggest that PIM1 upregulation is involved in lung cancer pathogenesis and is significantly associated with resistance to chemotherapy in NSCLC patients.

**Figure 1.**
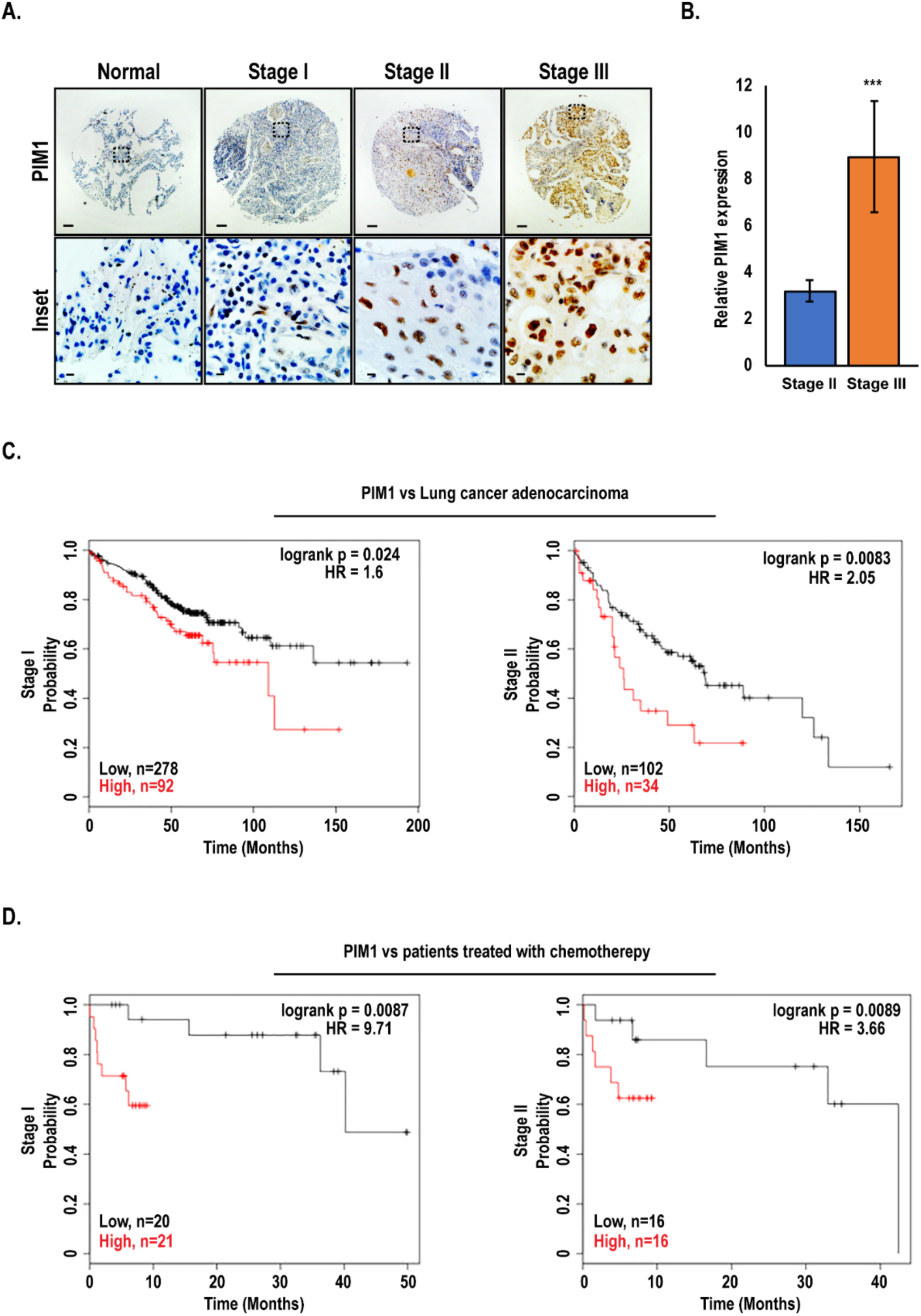
PIM1 is upregulated in advanced lung cancer and predicts poor survival outcomes: (A) Representative immunohistochemical staining of PIM1 expression by clinical stage in human lung cancer tissue array (main section, scale bars 100 μm; inset, scale bars 20 μm). (B) Quantification of average PIM1 expression in stage II vs. stage III of human lung cancer, mean ± SEM, n=37. ***, p<0.001. (C) Kaplan-Meier analysis of overall survival in human lung cancer adenocarcinoma patients with high vs. low PIM1, and (D) overall survival of patients treated with chemotherapy with high vs. low PIM1.

### PIM1 inhibition augments mitochondrial superoxide production and ROS accumulation

The fragmented or fused state of mitochondria is critical for maintaining proper function. One of the earliest signs of compromised mitochondria is amplified superoxide production, which ultimately leads to increased production of ROS. Because PIM inhibitors cause a dramatic increase in ROS, we hypothesized that PIM inhibition could generate excess ROS by impairing mitochondrial function. To test this, MitoSOX was used to selectively measure superoxide levels at the mitochondria in WT and Triple knockout MEFs (TKO; lacking all 3 PIM isoforms), and fold change in corrected total cell fluorescence (CTCF) intensity was measured. TKO MEFs had high basal superoxide levels compared to WT MEFs, and TKO MEFs with PIM1 added back (TKO-PIM1) displayed significantly reduced superoxide (Fig 2A). Similarly, treatment of WT MEFs with a pan-PIM kinase inhibitor (PIM447) caused a 2-fold amplification in superoxide production (Fig 2B). To validate that the observed effects are specific to inhibition of PIM and not an artifact of the drug itself, we treated a panel of NSCLC cell lines (H1299, A549, and H460) with a chemically distinct pan-PIM kinase inhibitor (AZD1208). A similar induction in superoxide production was observed in response to AZD1208 in all cell lines tested, indicating that these effects are specific to PIM inhibition (Fig 2C). Moreover, live cell imaging of mitoSOX demonstrated that superoxide levels were induced within 2 h of treatment with PIM inhibitors, indicating that blocking PIM has a direct effect on mitochondrial function, independent of its ability to inhibit the Nrf2 antioxidant pathway (Fig 2D and Fig S1). To further establish that compromised mitochondrial function following PIM inhibition ultimately causes free radical accumulation, we treated lung cancer cells with AZD1208 for 48 hours and measured ROS levels by EPR spectroscopy. Each of the NSCLC cell lines tested showed a dose-dependent increase in cellular ROS levels in response to PIM inhibition (Fig 2E). These results indicate that PIM1 loss or inhibition impairs mitochondrial function and leads to the accumulation of cellular ROS.

**Figure 2.**
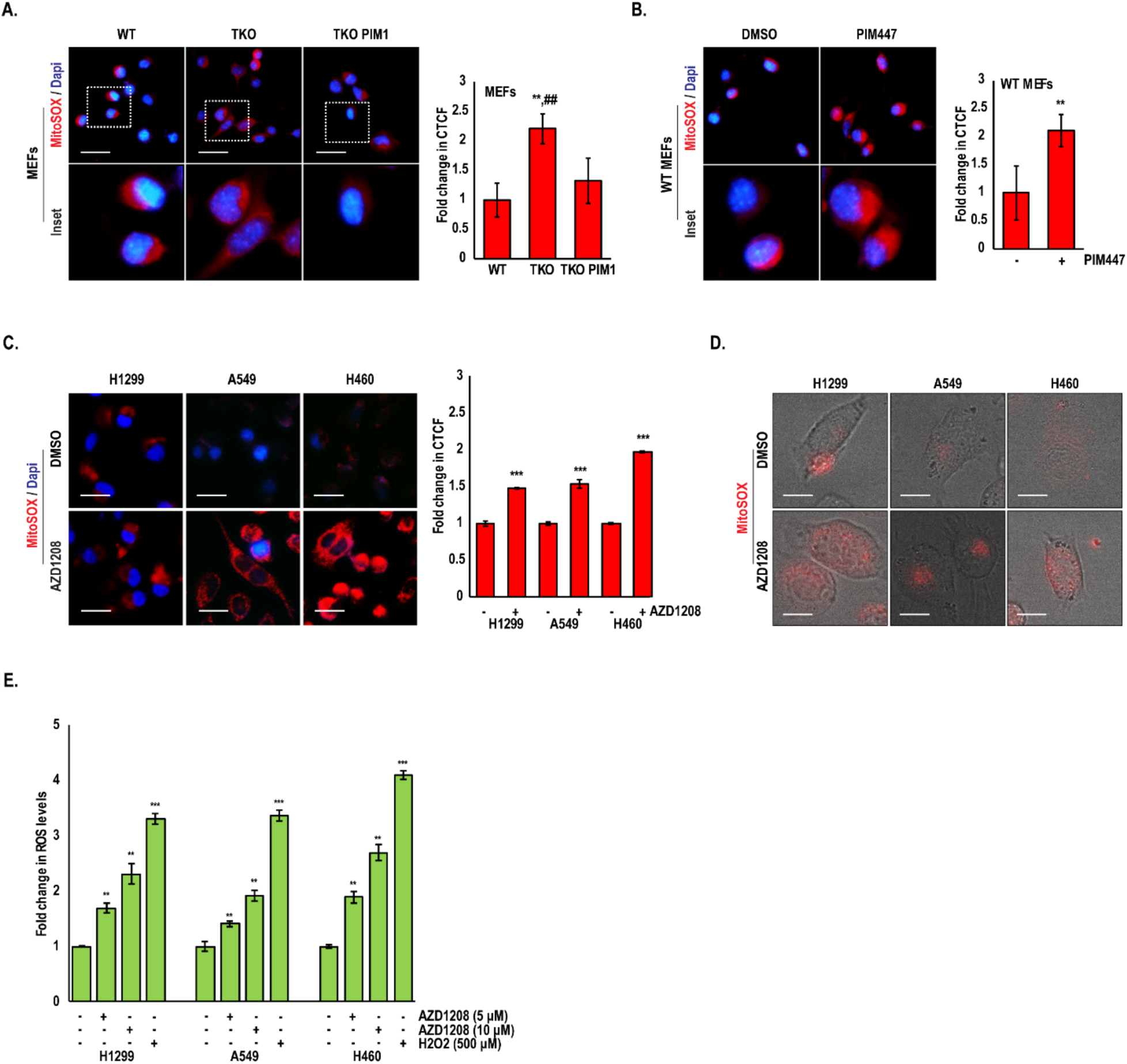
PIM inhibition increases mitochondrial superoxide production and total cellular ROS: (A) Mitochondrial ROS was detected by MitoSOX staining in WT, TKO, and TKO-PIM1 MEFs and (B) WT MEFs treated with PIM447 (3 μM) for 24 h. (C) The indicated lung cancer cells were treated with AZD1208 (5 μM) for 24 h, and (D) MitoSOX staining was quantified as the fold change in CTCF. Representative micrographs of live cell imaging of MitoSOX (red) staining in lung cancer cells treated with AZD1208 (5 μM) for 2 h. (E) Fold change in ROS levels as measured by EPR in lung cancer cells treated as indicated; ROS levels, nM/min/mg protein; mean ± SD of three independent experiments. **, p<0.01 vs WT MEFs; ^##^, p<0.01 vs TKO MEFs; *, p<0.05, **, p<0.01, ***, p<0.001 vs DMSO. Scale bars, 20 μm.

### PIM1 loss or inhibition induces mitochondrial fragmentation

Based on our finding that small molecule PIM inhibitors increase superoxide as a precursor to intercellular ROS, we speculated that manipulating PIM1 expression and activity could alter mitochondrial dynamics. To test this hypothesis, we assessed the mitochondrial phenotype of WT MEFs, TKO MEFs and TKO-PIM1. Cells were stained with a mitochondrial marker, TOM20, and visualized by fluorescence microscopy. Immunoblotting of IRS1 (S1101), an established PIM substrate (24), was used as a control to monitor PIM activity. The number of fragmented mitochondria (< 1 μm) in TKO MEFs was approximately 3-fold higher than that in WT MEFs (Fig 3A-C). Reintroduction of PIM1 to TKO MEFs restored mitochondria to a more elongated state (>1.5 μm), similar to that observed in WT MEFs (Fig 3A-C). Next, A549 cells were treated with PIM447 and mitochondrial morphology was quantified by immunofluorescence. Similar to the effect of PIM knockout, small molecule PIM inhibitors significantly increased mitochondrial fragmentation by 3 to 4-fold compared to control (Fig 3D-F). Parallel experiments using AZD1208 or PIM1 overexpression in prostate (Du145), breast (SUM159 and MCF7), and colon cancer (RKO) cell lines showed similar results, indicating that PIM regulates mitochondrial phenotype a broad range of cancers (Fig. S2). The changes in mitochondrial phenotype observed in response to PIM inhibition were further validated using transmission electron microscopy (TEM) in H1299 lung cancer cells treated with AZD1208. Quantification revealed a significant increase in fragmented mitochondria and a decrease in elongated or intermediate (1-1.5 μm) mitochondria in response to pharmacological inhibition of PIM (Fig 3G-I and Fig. S3A). To verify that the effects of AZD1208 on mitochondria were independent of Nrf2, a parallel experiment was performed in H1299 cells with stable knockdown of Keap1, which have constitutive activation of Nrf2. PIM inhibition did not alter Nrf2 levels (Fig S3B) in these cells but significantly increased fragmentation (Fig S3A). Taken together, these findings suggest that PIM controls mitochondrial dynamics in lung cancer.

**Figure 3.**
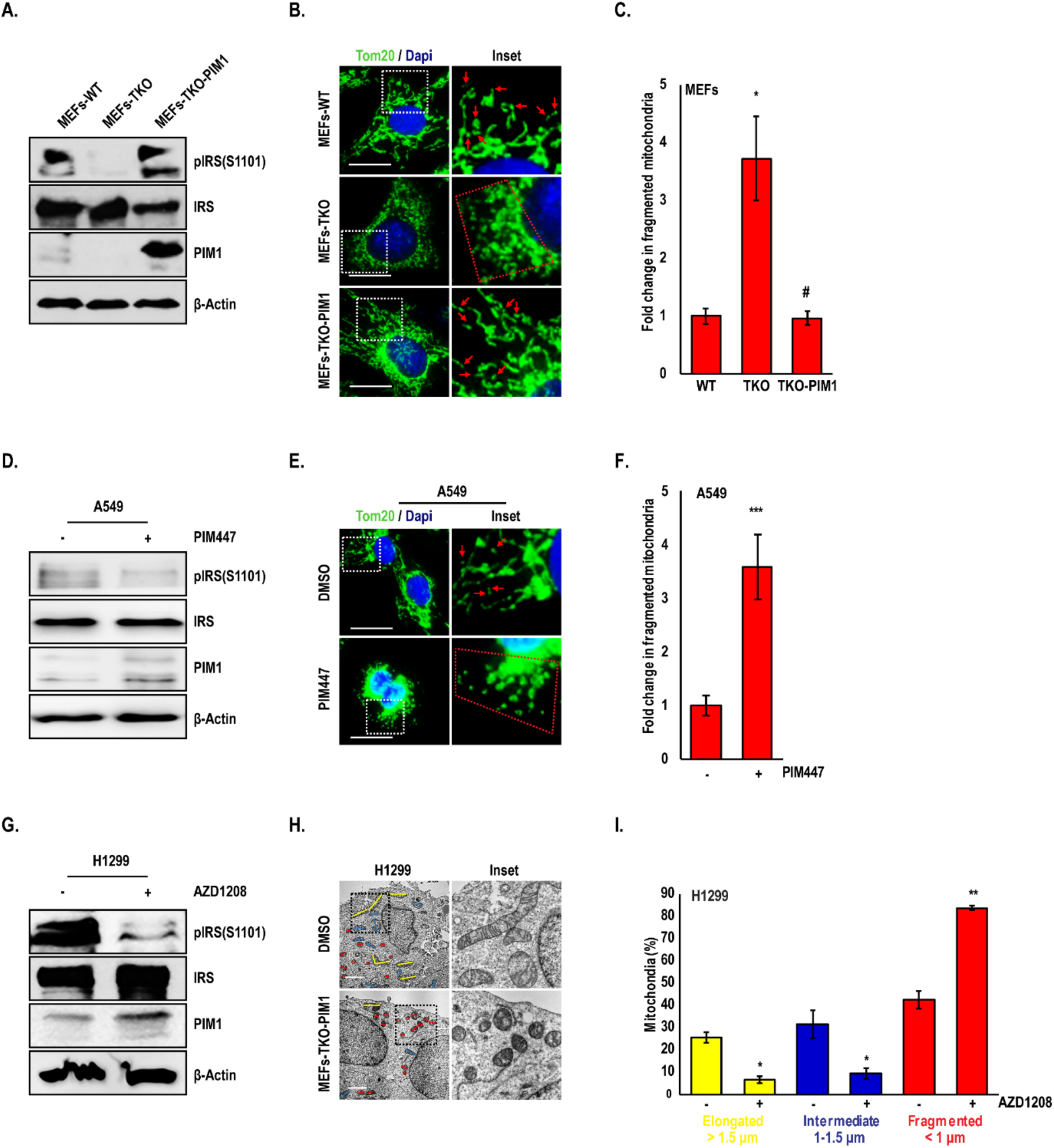
Loss of PIM induces mitochondrial fragmentation: (A) Representative western blots showing PIM levels and activity in MEFs. (B) Mitochondrial phenotype was assessed by TOM20 immunofluorescence (green), (C) and fragmented mitochondria (< 1 μm) were quantified (red arrowheads and mitochondria within red dotted line indicate fragmented mitochondria. (D-F) A549 cells were treated with PIM447 (3 μM) for 24 h and mitochondrial phenotype was assessed by TOM20 staining. (G) H1299 cells were treated with AZD1208 (5 μM) for 24 h, and (H) mitochondria were imaged by TEM. (I) Percentage of mitochondria with elongated (> 1.5 μm, yellow), intermediate (1-1.5 μm, blue) and fragmented (< 1 μm, red) mitochondria were quantified. Results are expressed as mean ± SD of three independent experiments. *, p<0.05 vs WT MEFs; #, p<0.05 vs TKO MEFs; *, p<0.05, **, p<0.01, ***, p<0.001 vs DMSO. Scale bars, 20 μm.

### PIM1 loss or inhibition affects mitochondrial phenotype in a Drp1-dependent manner

Drp1 is a key regulator of mitochondrial fission that promotes mitochondrial fragmentation when active. Therefore, we hypothesized that PIM could be controlling Drp1 to alter mitochondrial phenotype in lung cancer. To test whether there was an association between PIM1 and Drp1 in human samples, we stained a human lung cancer TMA for PIM1 and Drp1. Interestingly, areas within tissue cores expressing high levels of PIM1 correlated with low Drp1 expression and vice-versa (Fig 4A), and the inverse relationship between these proteins was further exaggerated during disease progression, as determined by assessing the PIM1/Drp1ratio by disease stage (Fig 4B). To verify the inverse correlation between these proteins, we immunoblotted for Drp1 in WT and TKO MEFs, as well as a panel of lung cancer cell lines (H1299, A549, and H460) treated with DMSO or PIM447. Both genetic depletion and chemical inhibition of PIM significantly increased total Drp1 levels (Fig 4C). Posttranslational modifications play an important role in maintaining Drp1 activity, particularly phosphorylation. Phosphorylation at S616 is thought to increase Drp1 activity and mitochondrial fragmentation, whereas phosphorylation at S637 opposes Drp1 activation and increases fusion. Thus, we next examined the phosphorylation state of Drp1 in response to PIM loss or inhibition. Strikingly, phosphorylation of S616 was significantly increased and phosphorylation of S637 was significantly reduced in TKO MEFs compared to WT MEFs (Fig. 4C). Similar effects on Drp1 phosphorylation were observed in lung cancer cells (H1299, A549, and H460) following treatment with PIM447 (Fig 4C). To confirm that Drp1 is responsible for the mitochondrial fragmentation observed in response to PIM loss or inhibition, TKO MEFs were treated with Mdivi-1, a Drp1 inhibitor, or incubated in HBSS as positive control to decrease fragmentation and induce fusion (25). After 24 h incubation with Mdivi-1, the basal number of fragmented mitochondria in TKO MEFs was significantly decreased (Fig 4D). Next, we tested whether Drp1 is necessary for PIM inhibitors to induce mitochondrial fragmentation. Treatment of A549 cells with PIM447 caused a 4-fold increase in fragmented mitochondria, whereas mitochondria were refractory to PIM inhibition when Drp1 was inactivated (Fig 4E). Because Mdivi-1 has been shown to have off target effects (26), we generated stable knockdown of Drp1 in A549 and H460 cells to verify our results using this inhibitor (Fig. 4F). Genetic suppression of Drp1 resulted in elongated/fused mitochondria and a significant decrease in fragmented mitochondria compared to the parental cell lines (Fig S4A). Importantly, mitochondria in cells lacking Drp1 were refractory to fragmentation (Fig 4G) and superoxide production (Fig S4B) upon treatment with PIM447. Thus, PIM inhibitors require the activation of Drp1 to induce mitochondrial fission and increase superoxide generation.

**Figure 4.**
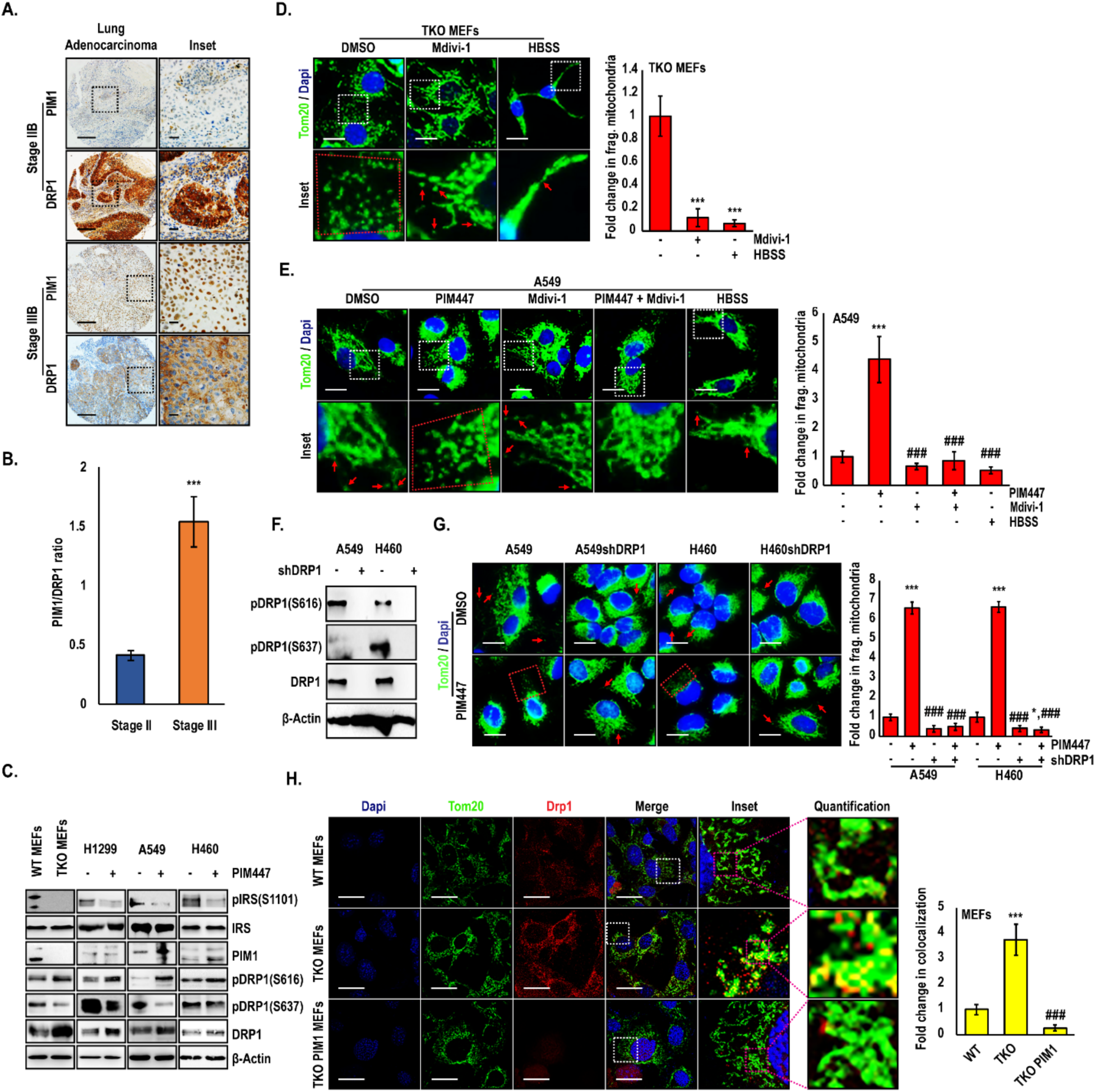
PIM1 affects mitochondrial phenotype in a Drp1-dependent manner: (A) Representative immunohistochemical staining and (B) quantification of PIM1 and Drp1 in NSCLC TMA (main section, scale bars 100 μm; inset, scale bars 20 μm). (C) Drp1 levels and phosphorylation were assessed by Immunoblotting in MEFs and lung cancer cell lines treated with DMSO or PIM447 (3 μM) for 24 h. (D) TKO MEFs were treated with a DRP1 inhibitor (Mdivi-1) and or HBSS and mitochondrial phenotype was assessed by TOM20 staining (green). (E) A549 cells were treated with the indicated drugs and mitochondrial phenotype was assessed by TOM20 staining (scale bars, 20 μm; ***, p<0.001 vs DMSO; and ^###^, p<0.001 vs PIM447). (F) Immunoblot and (G) Immunofluorescence analysis of TOM20 in Drp1 knockdown lung cancer cells treated with PIM447 (3 μM) for 24 h; Magnification 60X; scale bars, 20 μm;. *, p<0.05; ***, p<0.001 vs DMSO treated parental line; ^###^, p<0.001 vs PIM447-treated parental line. (H) WT, TKO, and TKO-PIM1 MEFs were stained for TOM20 (green) and Drp1 (red) and co-localization was quantified as the fold change in co-localized pixels (yellow); ***, p<0.01 vs WT MEFs; ^###^, p<0.01 vs TKO MEFs). Values are mean ± SD of 3 independent experiments. Scale bars, 20 μm.

Activation of Drp1 via phosphorylation is known to regulate its localization to mitochondria, so we reasoned that loss of PIM would increase the localization of DRP1 to mitochondria. To this end, we used confocal microscopy to assess the co-localization of Drp1 and mitochondria in WT and TKO MEFs. MEFs lacking PIM displayed a significant increase in TOM20 and Drp1 co-localization compared to WT MEFs (Fig 4H). Importantly, addback of PIM1 was sufficient to reduce total Drp1 levels as well as its localization to mitochondria, supporting the elongated phenotype observed in these cells (Fig 3B). Taken together, these results confirm that PIM loss or inhibition increases Drp1 levels and promotes its activity by altering the ratio of activating and inhibitory phosphorylation, ultimately increasing the recruitment of Drp1 to mitochondria, where it increases fission.

### PIM inhibition sensitizes lung cancer cells to chemotherapy by altering mitochondrial phenotype

Elevated PIM1 expression and elongated mitochondrial morphology have been linked to drug resistance, but the association between these factors has not been tested. We proposed that elongated mitochondrial phenotype associated with PIM1 upregulation is essential for PIM1 to promote chemoresistance in lung cancer. To validate this, A549 and H460 lung cancer cell lines were stably transfected with vector control (pCIP) or PIM1 (hPIM1) plasmids and treated with docetaxel for 72 h (Fig 5A and 5B). Immunoblotting demonstrated that PIM1-overexpressing cells had reduced total Drp1 levels and altered phosphorylation status (increased S637 and decreased S616). Indeed, cells overexpressing PIM1 displayed an elongated mitochondrial phenotype (Fig 5C) and were significantly resistant to docetaxel (IC_50_ values for A549 pCIP vs. hPIM1: 2.2 vs. 3.9 nM and H460: 1.7 vs. 3.4 nM) and cisplatin (IC_50_ values for A549 pCIP vs. hPIM1: 1.9 to 3.3 μM and H460: 1.5 vs. 4.4 μM) (Fig 5D). Since PIM1 levels are significantly higher in advanced lung cancer (Fig 1A), and its overexpression can lead to chemoresistance, we hypothesized that PIM inhibition would sensitize lung cancer cells to chemotherapy. To test this, A549, and H460 cells were treated with AZD1208 and docetaxel for 72 h and cell viability was measured by crystal violet staining. Dose-response curves and CI calculation revealed a synergistic anti-tumor response of combined treatment with AZD1208 and docetaxel (Fig S5A-C). To confirm that mitochondrial fragmentation is required for PIM inhibitors to sensitize lung cancer cells to chemotherapy, we treated shDRP1 cells with AZD1208 and docetaxel. Noticeably less synergy was observed between the PIM inhibitor and docetaxel in cells lacking Drp1, indicating that the ability of PIM inhibitors to cause fragmentation is critical for their ability to sensitize lung cancer cells to chemotherapy (Fig S5D-F). To determine whether combined treatment of PIM and docetaxel places added stress on mitochondria and enhances the accumulation of intercellular ROS, A549 and H460 cells were stained with H2DCF-DA and ROS was measured by flow cytometry. Inhibition of PIM alone amplified ROS in both A549 and H460 cells, and the combination of AZD1208 and docetaxel caused a further significant increase in ROS (Fig 5E).

**Figure 5.**
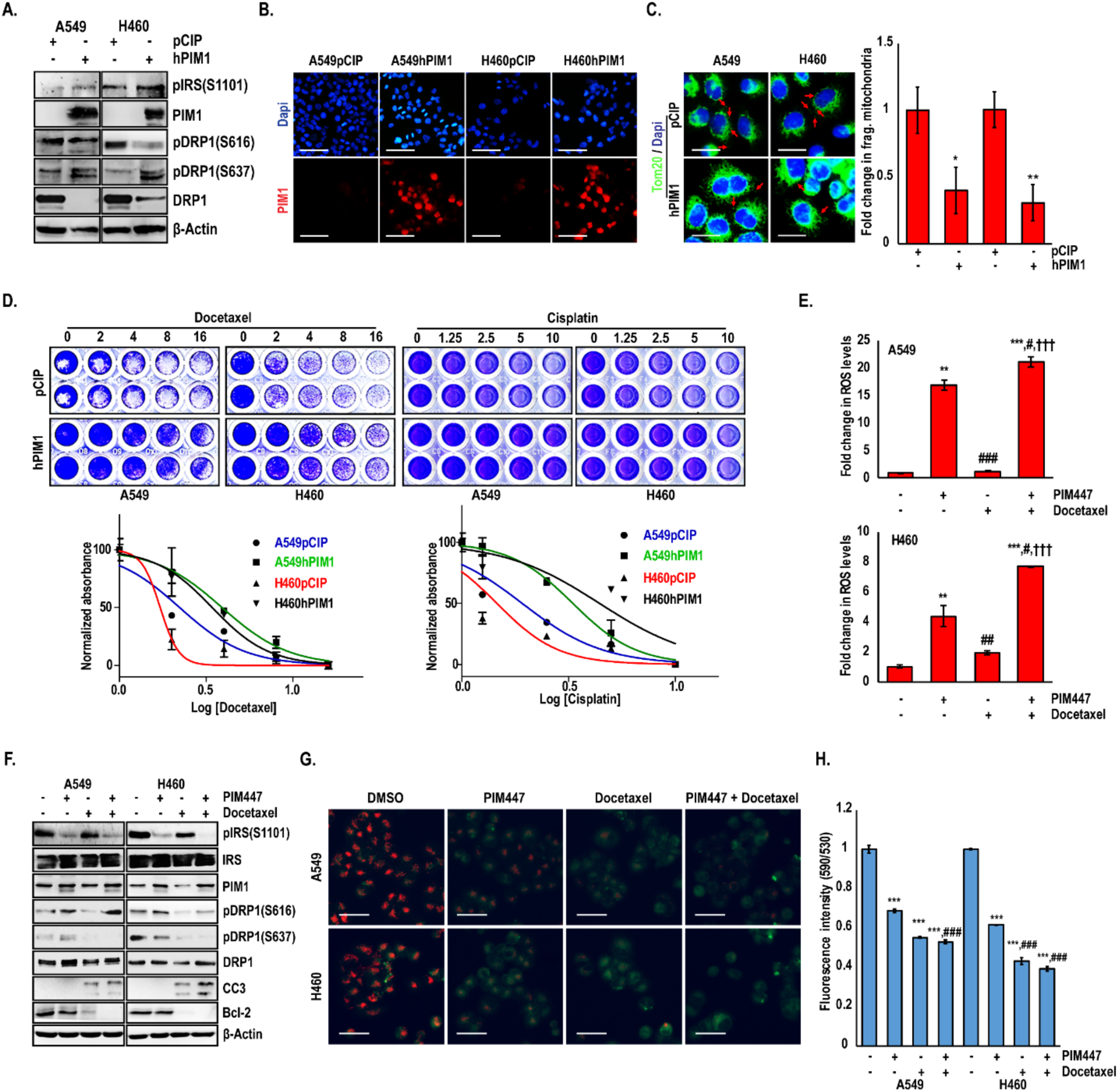
PIM inhibition sensitizes lung cancer cells to chemotherapy by altering the mitochondrial phenotype. Stable PIM1 overexpression in lung cancer cells confirmed by (A) western blotting and (B) Immunofluorescence analyses. (**C)** Mitochondrial phenotype assessed by TOM20 immunofluorescence (green) in A549 and H460 cells with PIM1 overexpression. Magnification 60X; scale bars, 20 μm. (*, p<0.05; **, p<0.01 vs pCIP transfected line). (D) Crystal violet cell viability assay and drug IC50 quantification in lung cancer cells after 72 h docetaxel or cisplatin treatment. (E) Flow cytometric analysis of H_2_-DCFDA intensity representing ROS levels in A549 and H460 cells treated with PIM447, docetaxel, or the combination. (F) Immunoblot analysis of A549 and H460 cells treated with PIM447 (3 μM), docetaxel (3 nM), or the combination. (G) Live cell micrographs of JC-1 aggregates in A549 and H460 cells treated with PIM447, docetaxel, or the combination (Scale bars, 40 μm), and (H) JC-1 staining was quantified using a fluorescence plate reader. Values are mean ± SD of 3 independent experiments. **, p<0.01, ***, p<0.001 vs DMSO; ^#^, p<0.05, ^##^, p<0.01, ^###^, p<0.001 vs PIM447; and ^†††^, p<0.001 vs Docetaxel.

To determine how PIM1 inhibition sensitizes lung cancer cells to docetaxel, we assessed markers related to mitochondrial phenotype and apoptotic cell death. Combination therapy induced the expression of markers of mitochondrial fragmentation and increased activation of the apoptotic pathway (i.e., downregulation of Bcl-2 and increased cleaved-caspase 3 (CC3)) (Fig 5F). To verify that cells treated with the combination were undergoing apoptosis initiated at the mitochondria, we measured the effect of combination therapy on changes in mitochondrial membrane potential (MMP), which is a precursor to cytochrome-c release and caspase activation, via JC-1 staining assay. A panel of lung cancer cells were first screened for JC-1 aggregates in the presence or absence of H_2_O_2_ to indicate changes in MMP, and a 24 h time point was chosen for subsequent drug treatment (Fig S5G). Cells treated with the combination of AZD1208 and docetaxel exhibited hyper depolarization of mitochondria that was greater than with either agent alone, indicative of the loss of MMP (Fig 5G and 5H).

### Combined treatment with PIM inhibitor and docetaxel has synergistic anti-tumor effects in a xenograft model of lung cancer

Based on our findings *in vitro*, we next investigated the effect of PIM1 inhibition and/or docetaxel treatment on tumor growth, proliferation, and apoptosis *in vivo*. Five million A549 cells were injected subcutaneously into each flank of SCID mice, and tumors were allowed to establish. Once the tumor size reached approximately 100-120 mm^3^, mice were randomly segregated into the following treatment groups: vehicle, AZD1208, docetaxel, and AZD1208 + docetaxel (Fig 6A). Tumor volume was measured by caliper every 3^rd^ day until sacrifice. The effect on tumor growth of either PIM inhibition or docetaxel alone was moderate, reducing overall tumor growth by approximately 50% compared to control. Strikingly, combination therapy produced a synergistic anti-tumor response that was significantly more effective than either agent alone (Fig 6B and 6C). The effect of each treatment regimen on tumor cell proliferation was assessed by immunohistochemical staining of Ki67. Although Ki67 staining was reduced in all treatment groups compared to vehicle, proliferation was reduced to the greatest extent in animals treated with AZD1208 and docetaxel (Fig 6D and 6E). Confirming our *in vitro* data that PIM inhibition induces apoptosis in docetaxel-treated cells, cleaved caspase 3 (CC3) was increased by nearly 4-fold with combination treatment compared to control or treatment with either agent alone (Fig 6D and 6E). To confirm that the signaling axis between PIM and Drp1 is intact and effectively altering the mitochondrial phenotype *in vivo*, we first assessed Drp1 levels in each of our treatment groups by immunohistochemistry. Treatment with docetaxel alone reduced Drp1 levels compared to vehicle, whereas AZD1208 significantly increased Drp1 protein levels by over 3-fold (Fig 6E). Importantly, combination therapy significantly increased Drp1 levels compared to AZD1208 alone. Next, tumor tissues (n = 3 tumors/group) were lysed, and immunoblotting was used to assess changes in Drp1 levels and phosphorylation status. PIM inhibition alone or in combination with docetaxel, increased Drp1 phosphorylation at S616 and decreased Drp1 phosphorylation at S637 (Fig 6F). Moreover, total levels of Drp1 and CC3 were significantly increased in combination-treated tumors. To confirm that PIM inhibition caused mitochondrial fragmentation *in vivo,* tumor tissues from control or AZD1208-treated mice were processed and imaged by TEM. Significantly more fragmented mitochondria were present in the AZD1208-treated tumors than the control tumors (Fig 6G). Quantification revealed that the average mitochondrial length in tumors was reduced from 1.1μm to 0.6 μm with PIM inhibition (Fig 6H). These findings indicate that, PIM inhibition sensitizes NSCLC tumors to chemotherapy via the induction of mitochondrial fragmentation, free radical generation, and apoptosis (Fig 6I).

**Figure 6.**
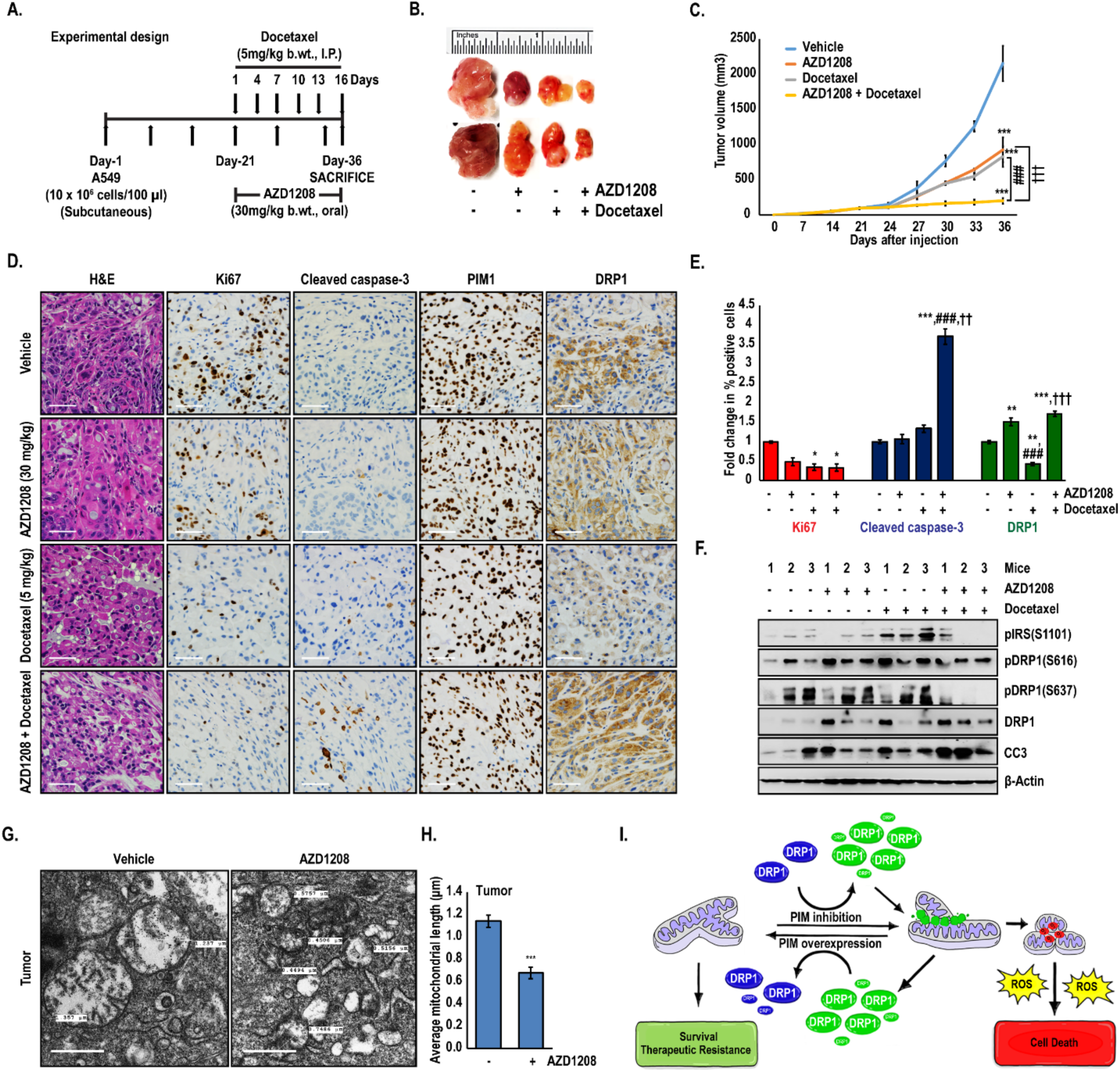
Combined treatment with PIM1 inhibitor and docetaxel displays synergistic anti-tumor effects *in vivo*. (A) Schematic of experimental design. (B) Representative tumors isolated from each treatment group (n=6). (C) Tumor volume (mm^3^) was determined over time (values ± SD; ***, p<0.001 vs vehicle; ^###^, p<0.001 vs AZD1208; and ^†††^, p<0.001 vs docetaxel). (D) Representative images of tumor sections (4 μm) examined by H&E and immunohistochemical analysis (scale bars, 20 μm). (E) Quantification of immunohistochemistry (*, p<0.05, **, p<0.01, ***, p<0.001 vs vehicle; ^###^, p<0.001 vs AZD1208; and ^††^, p<0.01, ^†††^, p<0.001 vs docetaxel). (F) Immunoblot analysis of tumor samples isolated from each group (n=3). (G) Tumors isolated from control and AZD1208-treated mice were imaged by TEM, and (H) mitochondrial length was quantified (scale bars, 500 nm) (n=49; values ± SEM; ***, p<0.001 vs vehicle). (I) Model describing the role of PIM in regulating the mitochondrial phenotype, ROS production and survival/therapeutic resistance.

## Discussion

Due to the limited efficacy of chemotherapy in NSCLC patients, the identification of new molecular targets to overcome drug resistance is central to improving patient prognosis. Emerging clinical and preclinical evidence demonstrates that the physical state of mitochondria significantly impacts the sensitivity of cancer cells to chemotherapy. Therefore, identifying anti-cancer agents that alter mitochondrial dynamics is a promising approach to overcome therapeutic resistance. Previous studies have shown that one mechanism by which cisplatin and docetaxel induce cell death by is by enhancing mitochondrial fragmentation and subsequent apoptosis (27,28). In the past, activation of PIM kinases has been implicated in chemoresistance via the phosphorylation of anti-apoptotic substrates, such as Bad and Bcl-2 (29,30). Here, we identify a new mechanism by which PIM1 contributes to chemoresistance in NSCLC: by promoting mitochondrial fusion.

The balance of mitochondrial fusion and fission affect mitochondrial function, which can impact many cellular processes involved in tumorigenesis, including changes in metabolism, proliferation, and apoptosis (31). However, the physiological consequences of shifting mitochondrial dynamics depend on the extent of fusion/fragmentation as well as environmental stress and genetic factors. For example, in RAS-driven pancreatic cancers, mitochondrial fission is required for proliferation, and knockdown of Drp1 is sufficient to block tumor growth, indicating a pro-tumorigenic role for fission (9,32). Alternatively, excessive mitochondrial fission in the same cell lines has been described to induce oxidative injury, deplete mitochondrial ATP, and induce apoptosis (33). Moreover, a fused mitochondrial network is correlated with cell survival and resistance to therapy in many types of cancer, including NSCLC (7,34). Because Drp1 is required for mitochondrial fission, understanding the cellular inputs that control its activation in cancer is critical for (S616: activating and S637: inhibitory) controls its levels and localization to mitochondria. sensing cellular energy stress: AMPK-AKAP-PKA (36,37). Interestingly, while the overall trend is consistently toward activation of Drp1, the relative impact of PIM inhibition on total levels of Drp1 and regulation of its phosphorylation status appears to differ in cell lines from different types of cancer. Inactivation of PIM significantly altered phosphorylation at both sites, increasing the activating phosphorylation at S616 and decreasing the inhibitory phosphorylation at S637. PIM1 overexpression was previously shown to increase phospho-Drp1 (S637) in cardiomyocytes isolated from transgenic mice (38), and our data corroborated these results in cancer cells. Thus, Drp1 is likely a direct target of PIM at the inhibitory S637 site, which aligns with the protective role of PIM in cancer. We also observed a marked increase in phospho-Drp1 (S616), which is typically associated with ERK/CDK1activation. Indeed, Inhibition of PIM significantly upregulated ERK phosphorylation compared to controls in a panel of lung cancer cell lines, whereas overexpression of PIM1 decreased ERK phosphorylation (Fig. S6). This finding is interesting, as crosstalk between PIM and ERK has not been previously described and warrants further investigation.

Our data suggest that PIM1 expression is particularly relevant to the dysregulation of apoptosis in NSCLC. PIM1 increases with disease progression in NSCLC and high expression of PIM1 is significantly associated with worse overall survival. PIM kinases are induced in response to cellular stress, including mechanical stress, nutrient deprivation, and hypoxia (20,39,40), so it is likely that changes in the tumor microenvironment account for the observed increase in PIM1 in later stage NSCLC. Supporting prior studies, we observe a significant increase in PIM1 levels following treatment with docetaxel *in vitro* and *in vivo*, providing further evidence of its role as a key player in the cellular response to stress (41). *In vitro*, overexpression of PIM1 decreased the sensitivity of NSCLC cell lines to docetaxel and cisplatin. This finding is of particular significance in the context of therapeutic resistance, as NSCLC patients with high PIM1 display a significantly worse response to chemotherapy than patients with low PIM1 (Fig 1B). Here, we demonstrate that PIM kinase inhibitors cause excessive mitochondrial fragmentation and triggered superoxide overloading, leading to oxidative stress and reduction in the MMP. These alterations were accompanied by an upregulation of proapoptotic proteins and a downregulation of antiapoptotic factors. Combined treatment with PIM inhibitors and docetaxel or cisplatin produced a synergistic cytotoxic response in NSCLC *in vitro* and *in vivo*. Moreover, PIM inhibitors sensitized NSCLC cell lines to chemotherapy in NSCLC cell lines with WT Ras (NCI-H2228) and mutant EGFR (NCI-H1975), indicating that this therapeutic strategy irrespective of mutational status (Fig S7). Importantly, the dependent on mitochondrial fragmentation, as stable knockdown of Drp1 in NSCLC cells blocked the ability of PIM inhibitors to induce mitochondrial fragmentation and increase ROS. In these cells, combining PIM inhibitors with chemotherapy was far less effective, demonstrating that the impact of PIM on the mitochondrial phenotype is central to its utility as a therapeutic target. It remains to be determined whether PIM also controls mitochondrial dynamics in primary cells, and if so, whether this would limit the therapeutic window for PIM inhibitors. Previous studies have described a role for PIM in regulating mitochondrial dynamics in cardiomyocytes (38), so it is likely that PIM is a general mediator of mitochondrial phenotype. However, PIM1 is upregulated in cancer and by many aspects of the tumor microenvironment, such as hypoxia. Therefore, we predict that the effect of PIM inhibitors on mitochondrial dynamics and ROS production would be more dramatic in cancer cells compared to primary cells, providing some specificity toward tumor tissue.

Tumors are constantly exposed to environmental stress that result in ATP depletion and increased oxidative stress. In order to adapt and survive, it is essential for tumor cells to maintain redox homeostasis. Previous work from our lab and others demonstrated that loss of PIM kinases reduces Nrf2 activity and lowers the antioxidant capacity of cancer cells, leading to the accumulation of ROS (42,43). Here, we describe a new role for PIM kinases in controlling redox homeostasis in cancer by controlling mitochondrial dynamics and increasing the production of ROS, independent of their ability to alter Nrf2 and the anti-oxidant response. Based on these findings, we propose that PIM serves a dual role in controlling oxidative stress. First, PIM maintains an elongated mitochondrial network, limiting acute ROS production at the mitochondria. At the same time, increased PIM stabilizes Nrf2 to increase the antioxidant capacity of the cell. Thus, inhibition of PIM both stimulates the production of ROS at the mitochondria by causing excessive fission and also decreases the antioxidant response by reducing Nrf2 activation. Together, these effects place cancer cells into a state of severe oxidative stress and render them hypersensitive to anti-cancer therapy.

## Materials and Methods

### Plasmids

pCIP and hPIM1 constructs were created by subcloning into the expression vector pCIG3 (pCMV-IRES-GFP, a gift from Dr. Felicia Goodrum, Addgene, plasmid #78264), modified to replace the GFP cassette with puromycin resistance gene. pSuper-Retro-puro-human shDrp1 was previously described (9).

### Reagents and antibodies

AZD1208 was acquired from AdooQ Biosciences. Docetaxel and Mdivi-1 were obtained from Selleck Chemicals. JC-1, mitoSOX-red and H_2_-DCFDA were purchased from Invitrogen. Primary antibodies for immunoblot and immunofluorescence analyses were purchased from BD Biosciences [Actin, 612656], Santa Cruz [Drp1, sc-271583, PIM1, sc-13513 and Bcl-2, sc-7382], and Cell Signaling Technology [TOM20, 4240S; pIRS (S1101), 2385S; PIM1, 3247S; pDRP1 (S616), 3455S; pDRP1 (S637), 4867S; and cleaved caspase-3, 9661S. All other materials and chemicals were of reagent grade.

### Cell transfection and immunoblotting

Wild type (WT) mouse embryonic fibroblasts (MEFs), triple-knockout (TKO; Pim1^−/−^, Pim2^−/−^, and Pim3^−/−^) MEFs (21), TKO MEFs stably expressing PIM1 (TKO-PIM1) (22), H1299, H1299Keap1^−/-^, A549, and H460 cells were maintained in DMEM containing 10% FBS and 1% penicillin/streptomycin. All cell lines were maintained at 37°C in 5% CO_2_ and were authenticated by short tandem repeat DNA profiling performed by the University of Arizona Genetics Core Facility. The cell lines were used for fewer than 50 passages and routinely tested for mycoplasma contamination. Stable retroviral transfections were carried out to generate PIM1 overexpressing (hPIM1) and Drp1 knockout (shDrp1) cells. Immunoblotting was performed as described previously (23).

### Mitochondrial superoxide assay

MitoSOX red staining was performed to measure mitochondrial superoxide production. Cells were plated in 6-well plates containing microscope coverslips and treated as indicated. Post-treatment, cells were incubated with 5 µM MitoSOX red dissolved in Hank’s Balanced Salt Solution (HBSS) for 30 minutes. Then, cells were washed with 1X DPBS and associated staining was determined either on mounted slides or in live cells using fluorescent microscopy.

### ROS assay

Electron paramagnetic resonance (EPR) was used to measure ROS in H1299, A549, and H460 cells treated with PIM447 for 48 h. Post-treatment, cells were incubated in Krebs-HEPES buffer, pH 7.4 containing 200 µM CMH (1-hydroxy-3-methoxycarbonyl-2,2,5,5-tetramethylpyrrolidine) probe for 30 minutes at 37°C. Cells treated with 500 µM hydrogen peroxide for the same incubation period served as a positive control. Fifty microliters of the Incubation solution were added to a glass EPR capillary tube (Noxygen Science Transfer & Diagnostics, Elzach, Germany) that was placed inside the cavity of the E-scan spectrometer for data acquisition. The parameter settings for acquisition were as follows: center field 1.99 g, microwave power 1 mW, modulation amplitude 9 G, sweep time 10 seconds, number of scans 10, and field sweep 60 G. Sample temperature was stabilized and kept at 37°C by the Temperature & Gas Controller “Bio III” unit, interfaced to the spectrometer. Spectra were recorded and analyzed by using Win EPR software (2.11 version). ROS was also measured by DCF staining, as described previously (20).

### Immunofluorescence

Cells were plated in 6-well plates containing microscope coverslips and treated as indicated. Post-treatment, cells were fixed with 10% buffered formalin for 20 min and kept in blocking solution (5% NGS and 0.3% TritonX-100 in PBS) for 60 min. Then, cells were incubated with anti-TOM20 (rabbit mAb, 1:500 dilution), or anti-PIM1 (mouse mAb, 1: 100 dilution) or anti-DRP1 (mouse mAb, 1:200 dilution) antibodies for 60 min. Following primary antibody incubation, cells were washed with 1× PBS and incubated in secondary antibodies (Alexa Fluor 568 goat anti-mouse and Alexa Fluor 488 goat anti-rabbit, 1:500 dilution) for 60 min. Finally, cells were mounted on glass slides with mounting media (Cell Signaling Technology 8961S, Prolong® Gold) containing DAPI. Images were taken at 60× magnification using a fluorescent microscope.

### Transmission electron microscopy

Cells or tumor tissue samples were fixed with 2.5% glutaraldehyde in 0.1 M PIPES buffer, pH 7.4 overnight at 4°C. The samples were then washed with 0.1M PIPES, pH 7.4 three times for 10 minutes each. The samples were then post-fixed with 1% osmium tetroxide in PIPES, pH7.4 for 1 h, washed with deionized water two times for 10 min, followed by 20 min in aqueous 2% uranyl acetate, and washed again with deionized water for 10 min. The samples were then dehydrated with a graded series of increasing concentrations of ethanol (50%, 70%, 90%, and 100%) in Pelco Biowave Pro microwave, set at 250W, 20°C, and vacuum for 40 seconds. The samples were then infiltrated (microwave, 1:1 Spurr’s resin ethanol, 250W, 20°C, vacuum 3 minutes and Spurr’s resin, 25W, 20°C vacuum twice three minutes each) and embedded in Spurr’s resin overnight at 60^0^°C. Ultrathin (60nm) sections were cut onto uncoated copper mesh grids and stained with 2% lead acetate for 2 min. The samples were examined using FEI CM12 transmission electron microscope operated at 80kV. Digital images were obtained in 8-bit TIFF format using a 4×4 digital camera.

### Cell viability assay and drug interaction assessment

The cell viability was measured by crystal violet staining. Briefly, cells were plated in 96-well plates, treated with inhibitor/drugs for 72 h, fixed in 4% formaldehyde, and stained with 0.1% crystal violet. The cells were lysed in a 1% sodium dodecyl sulfate solution, and absorbance was measured using microplate reader at a wavelength of 595 nm. Synergy between PIM inhibitor and chemotherapies was assessed by calculating the combination index (CI) value using CompuSyn software. CI < 1 indicated synergy (the smaller the value, the greater the degree of synergy), CI = 1 indicated an additive effect, and CI > 1 indicated antagonism.

### JC-1 assay

Mitochondrial membrane potential (MMP) was determined by JC-1 assay. A549 and H460 cells were plated in 96-well plates and allowed to adhere overnight. Then, the cells were washed once with 1× DPBS, stained, and treated as indicated. At the end of the treatment period, absorbance was recorded using a fluorescence microplate reader with an excitation of 485 nm and emissions of 540 nm and 590 nm. Hydrogen peroxide was used as a positive control. MMP was determined using the ratio of the fluorescence of J-aggregates (590 nm) to monomers (540 nm) and normalized to the respective DMSO control. For JC-1 imaging, cells were seeded on 6-well plate, treated as indicated and processed as described above. Finally, images were taken for green and red channels at 20× magnification using a fluorescent microscope.

### *In vivo* studies

The sample size justification of 4 mice (8 tumors) per group is based on comparing the combination to each individual agent alone. We will have 80% statistical power to detect a standardized decrease of 1.325 between groups (difference divided by standard deviation) assuming a two-sided alpha of 0.05. Five million A549 cells in PBS were injected subcutaneously into each flank of SCID mice in PBS. Once average tumor size reached approximately 100 mm^3^, mice were randomized for treatment with vehicle (Cremophore EL/Ethanol/PBS-24/6/70 ratio, p.o. daily; 5% DMSO + 30% PEG + 5% Tween80 + ddH_2_O, i.p. every 3^rd^ day), AZD1208 (30mg/kg by p.o. daily), docetaxel (5 mg/kg by i.p. every 3^rd^ day), or AZD1208 + docetaxel. Tumor volume was monitored by caliper measurements. Thirty-six days after injection, animals were sacrificed, and tumors were harvested. Tumors were fixed, embedded in paraffin, and sectioned for staining with hematoxylin and eosin (H&E) or antibodies specific for Ki67, Cleaved caspase-3 (CC3), PIM1, and DRP1. Percent positive staining for the above mentioned proteins was calculated using ImageJ analysis software. Investigators were blinded to the sample information prior to software-based analysis. For Transmission electron microscopy (TEM) analysis, tumors from vehicle and AZD1208-treated mice were fixed in 2.5% glutaraldehyde. All animal studies were approved by the Institutional Animal Care and Use Committee at the University of Arizona.

### Statistical analysis

Tumor growth was analyzed by fitting a mixed linear model of tumor volume vs. time for each mouse. The resulting slopes (growth rate) was compared using a factorial model with vehicle, AZD alone, Docetaxel alone, and the AZD + Docetaxel dual treatment. Differences in proliferation and apoptosis among treatment groups was analyzed using linear mixed models adjusted for the correlation among measurements within the same mouse. All immunofluorescence staining and western blots are representative of at least three independent experiments. Differences across groups were determined by unpaired 2-tailed Student’s t-test. One-way analysis of variance (ANOVA) was used to analyze differences between more than two groups across one timepoint. P values were adjusted using Bonferroni’s multiple comparison test. The data is presented as the mean SD or mean SEM as indicated, and a p-value < 0.05 was considered statistically significant.

## Acknowledgements

We would like to thank Dr. Donna Zhang (University of Arizona) for providing the H1299 Keap1^−/-^ cell line and assistance with the acquisition and analysis of EPR results. We thank Adam R. Kohr for his assistance with graphic design. The research was supported by American Cancer Society grant RSG-16-159-01-CDD, American Lung Association grant LCD-504131, and Department of Defense PCRP Award (W81XWH-19-1-0455) to NAW. Cancer Center Support Grant P30CA023074 also provided support for this research.

## Role in the study

Study concept and design: NAW, SSC

Acquisition of data: SSC, RKT, CCJ, ALC, NAW

Analysis and presentation of data: NAW, SSC

Material support: DFA

Study supervision: NAW

Funding: NAW

---

**Figure S1:**
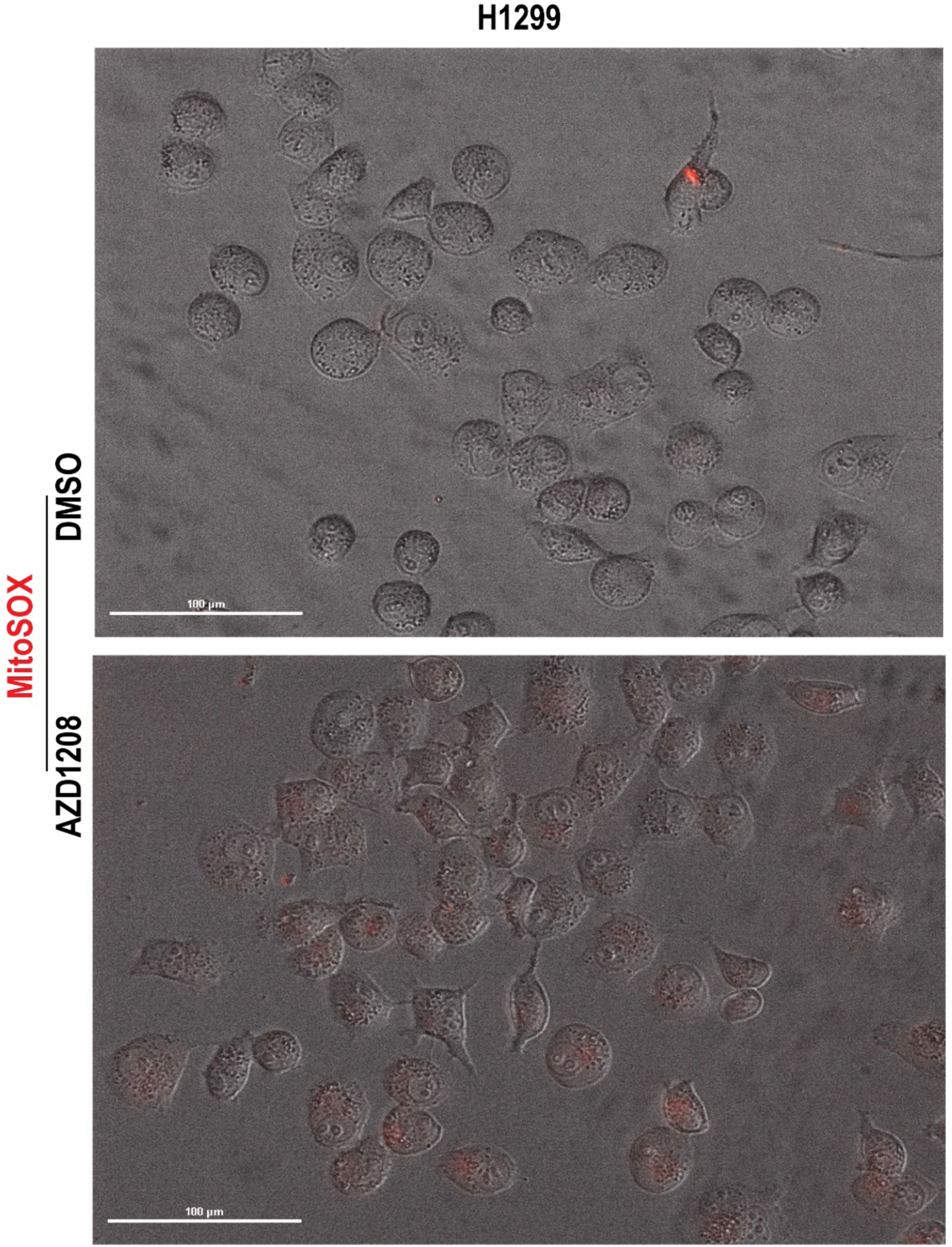
Live cell imaging showing increased MitoSOX staining (red) or superoxide production in H1299 cells treated with AZD1208 (5 μM) for 2h. Scale bars, 100 μm.

**Figure S2:**
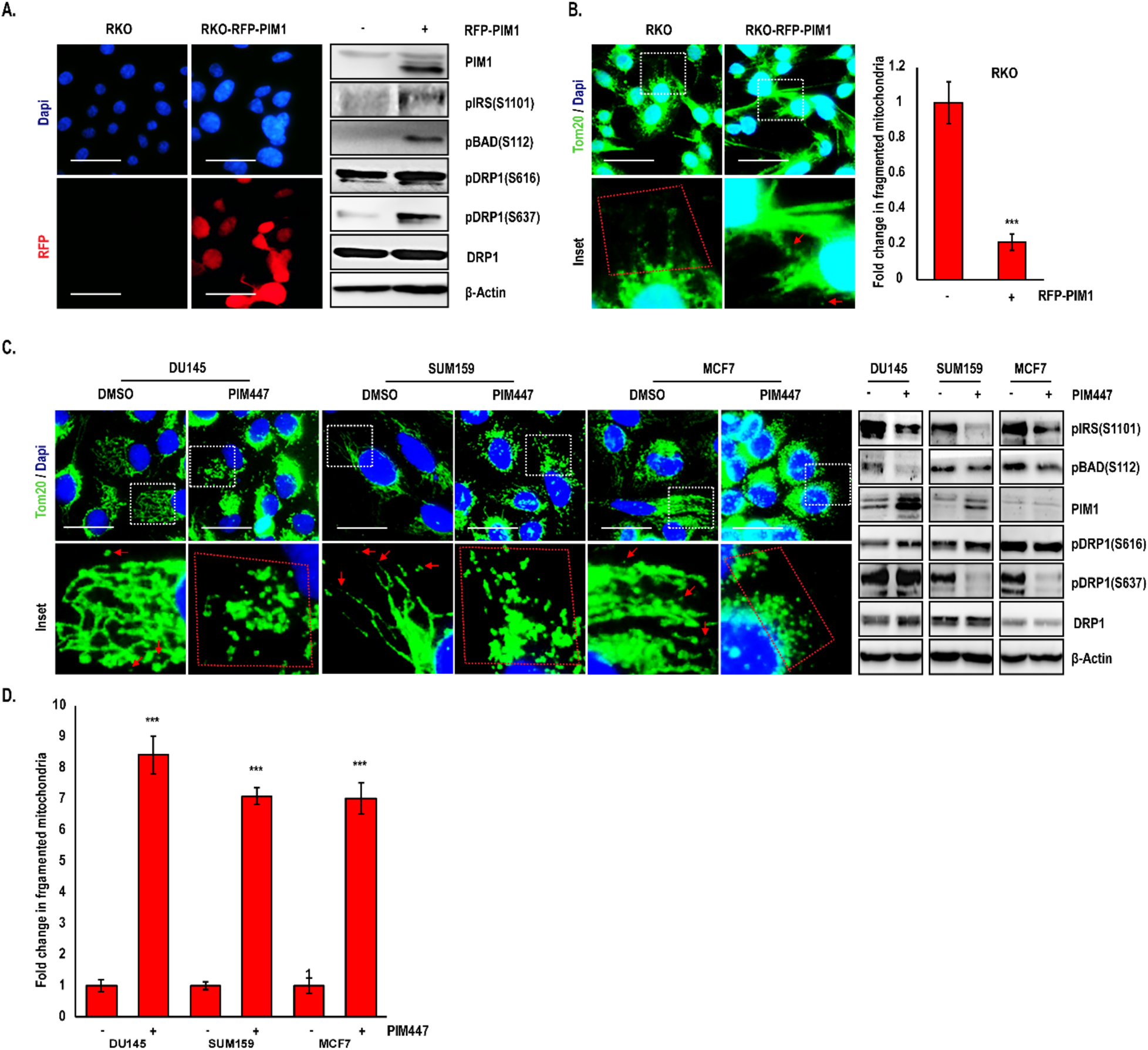
(A) Representative immunofluorescence images showing RFP-PIM1 transfection in RKO (colon cancer) cells. Western blot confirming increased DRP1 phosphorylation at S637 upon PIM1 overexpression. (B) Indicated colon cancer cells stained for TOM20 showing decrease in fragmented mitochondria with PIM1 overexpression. (C) Prostate (DU145) and breast (SUM159 and MCF7) cancer cells showing increased mitochondrial fragmentation (TOM20 staining) and altered DRP1 levels and/or phosphorylation with PIM447 treatment. (D) Fold change in fragmented mitochondria in indicated cells treated with PIM447. Red arrowheads and mitochondria within red dotted line indicate fragmented mitochondria (B & C). Results are expressed as mean ± SD of three independent experiments. ***, p<0.001 vs RKO; vs DMSO. Scale bars, 20 μm.

**Figure S3:**
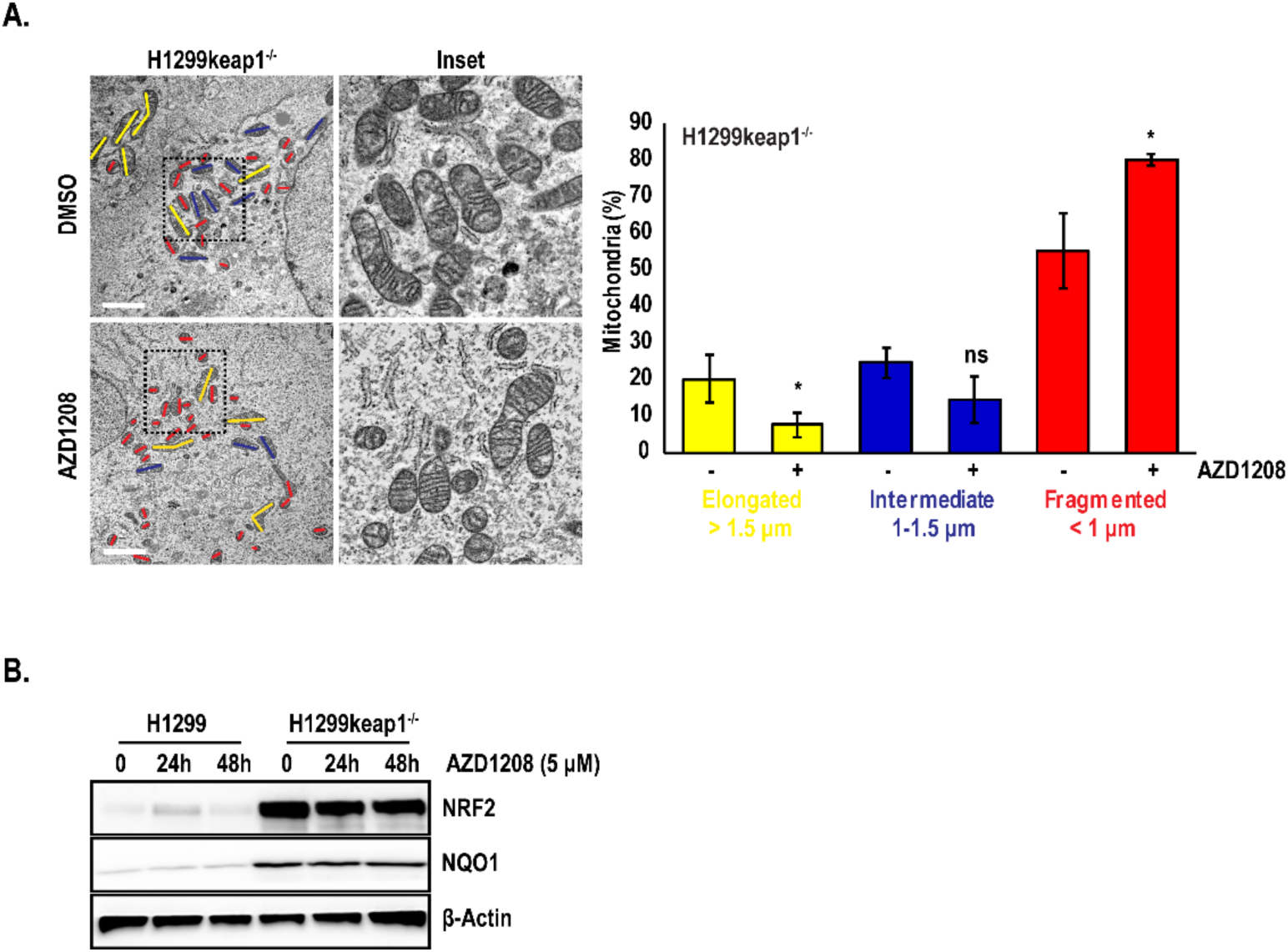
(A) TEM analysis showing increased mitochondrial fragmentation in H1299Keap1^−/-^ cells treated with AZD1208 (5 μM) for 24h. Scale bars, 2 μm. Histogram represents quantification of TEM images showing a significant change in the proportions of elongated (> 1.5 μm, yellow), intermediate (1-1.5 μm, blue) and fragmented (< 1 μm, red) mitochondria. (B) Immunoblotting in H1299 and H1299Keap1^−/-^ cells treated with AZD1208 (5 μM) for 0, 24h and 48h. Results are expressed as mean ± SD of three independent experiments. *, p<0.05 vs DMSO.

**Figure S4:**
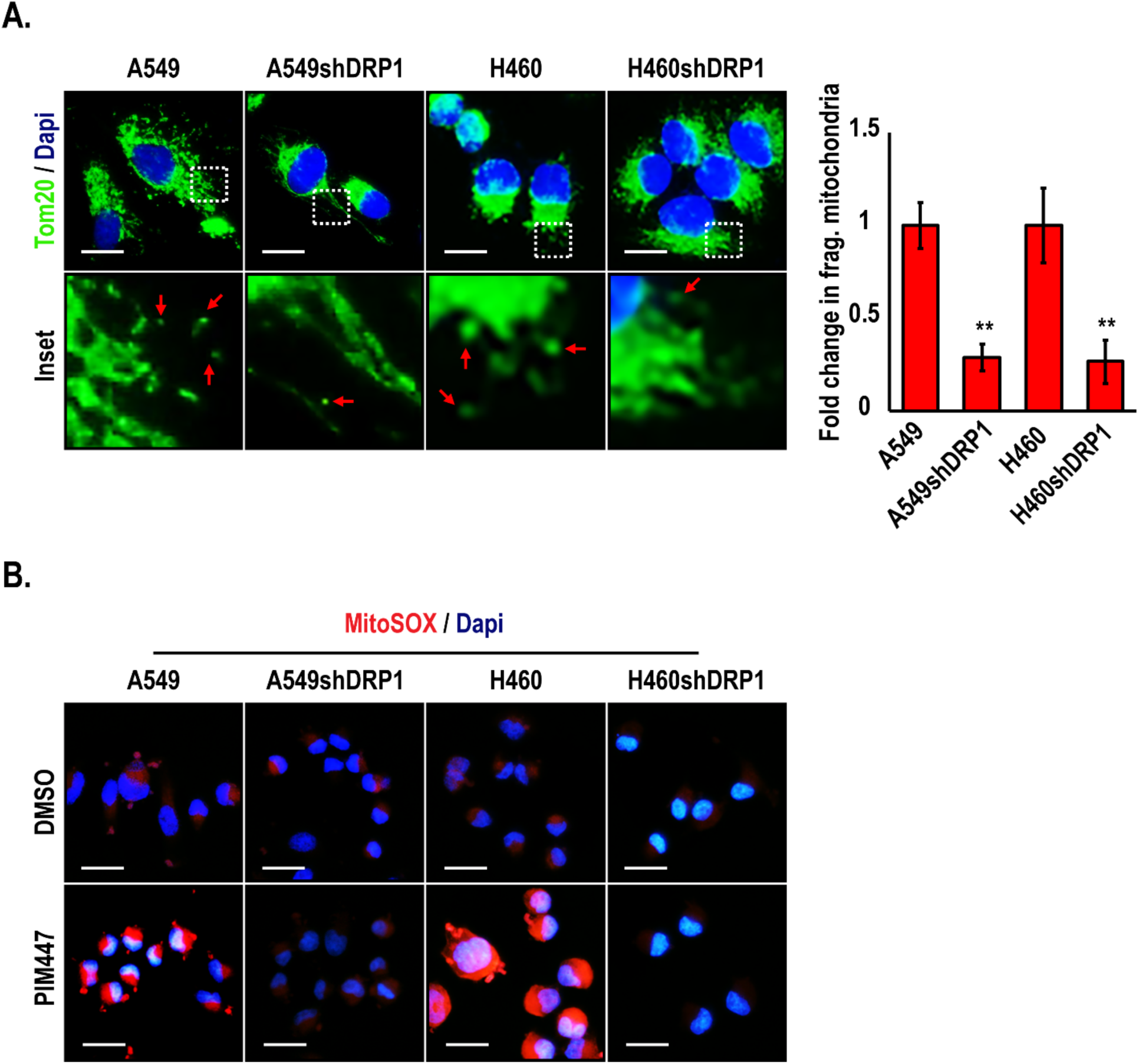
(A) Immunofluorescence analysis of TOM20 in Drp1-knockdown lung cancer cells (A549shDRP1 and H460shDRP1). Magnification 60X; scale bars, 20 μm; red arrowheads indicate fragmented mitochondria (< 1 μm). Histogram representing fold change in the proportion of fragmented mitochondria. Results are expressed as mean ± SD of three independent experiments. **, p<0.01 vs parental line. (B) MitoSOX (red) staining in Drp1 knockout lung cancer cells treated with PIM447 (3 μM) for 2 h. Results are expressed as mean ± SD of three independent experiments.

**Figure S5:**
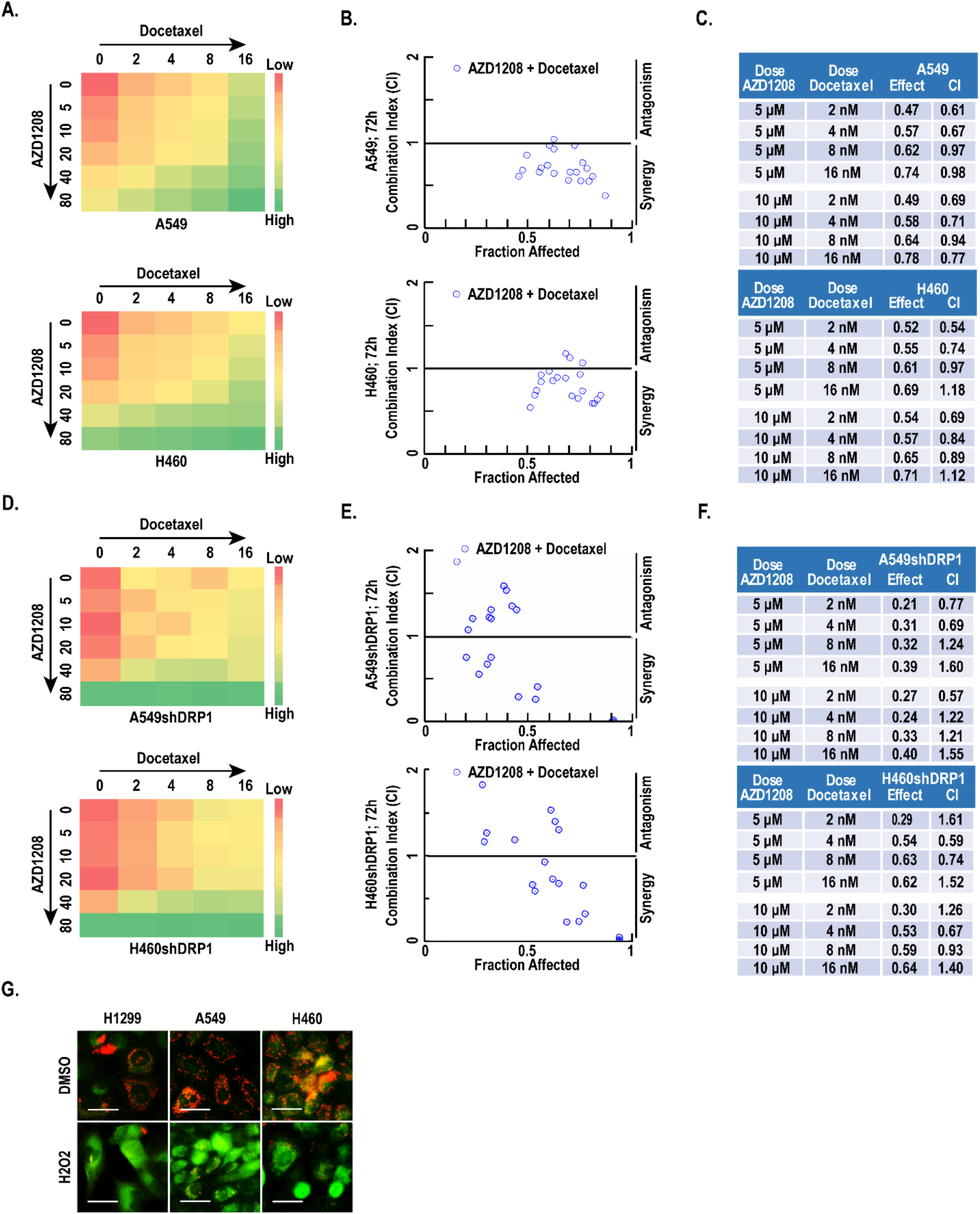
(A) Heat map representing dose-response, (B) combination index (CI) representing synergy, and (C) table showing CI values in A549 and H460 cells cotreated with a range of AZD1208 (μM) and docetaxel (nM) for 72 h. (D) Heat map representing dose-response, (E) combination index (CI) representing synergy, and (F) table showing CI values in A549shDRP1 and H460shDRP1 cells cotreated with a range of AZD1208 (μM) and docetaxel (nM) for 72 h. (G) JC-1 staining in lung cancer panel with H_2_O_2_ treatment for 24h. Magnification 40X; scale bars, 20 μm. Results are expressed as mean ± SD of three independent experiments.

**Figure S6:**
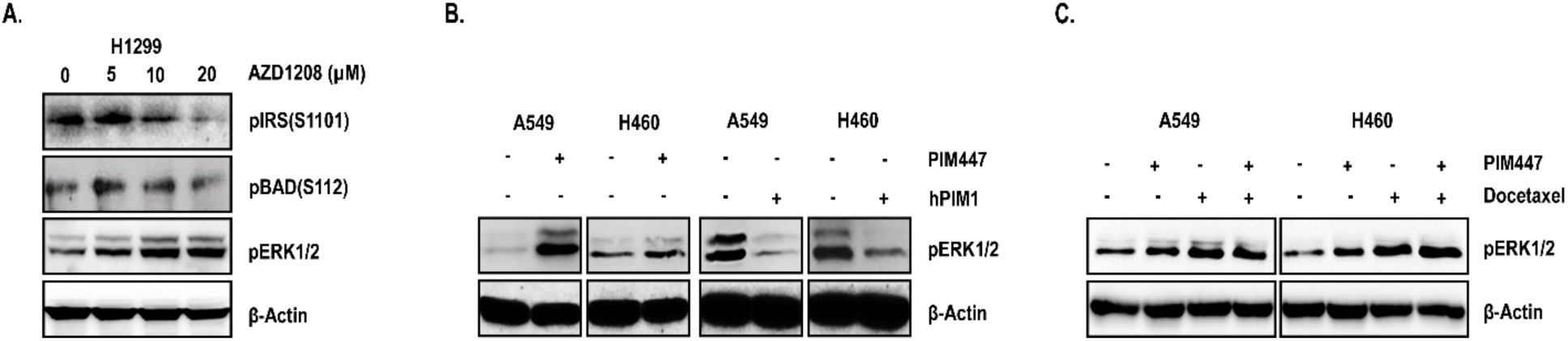
Immunoblot analysis of indicated cells treated with (A) μM concentration of AZD1208, (B) PIM1 overexpression plasmid (hPIM1) and PIM inhibitor (PIM447, 3 μM), (C) PIM447 (3 μM), docetaxel (3 nM), or the combination for 24h.

**Figure S7:**
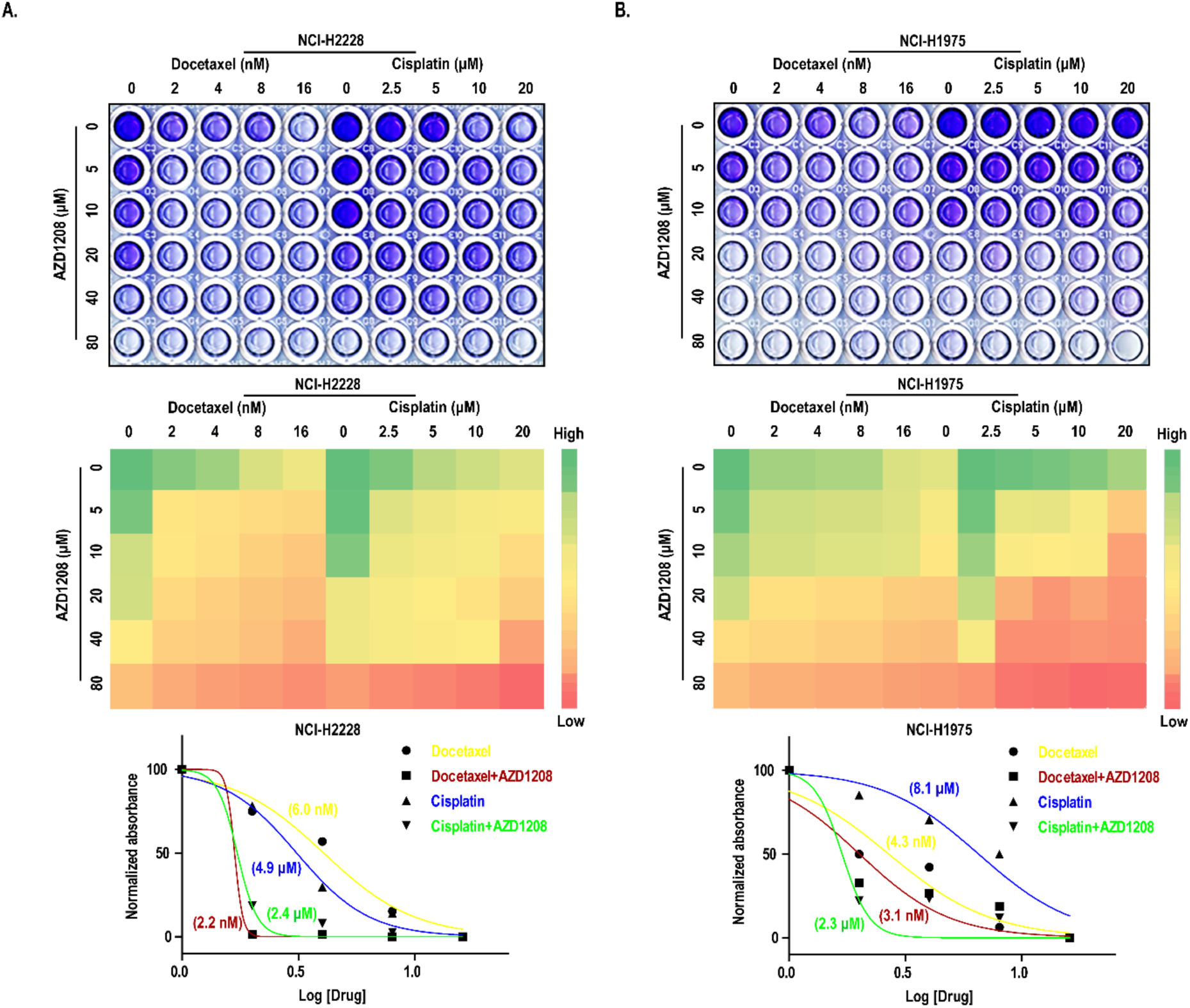
Crystal violet cell viability assay, heat maps representing cell viability, and drug IC_50_quantification in (A) wild type RAS (NCI-H2228) and (B) EGFR mutated (NCI-H1975) lung cancer cells after 72 h docetaxel or cisplatin treatment, alone, and in combination with AZD1208. Decrease in docetaxel or cisplatin IC_50_ values AZD1208 at 5 μM concentration indicated cell sensitization.

